# Gel-Assisted Proteome Position Integral Shift (GAPPIS) Assay Returns Molecular Weight to Shotgun Proteomics and Identifies Novel Caspase 3 Substrates

**DOI:** 10.1101/2024.02.14.579877

**Authors:** Zhaowei Meng, Amir Ata Saei, Xuepei Zhang, Hezheng Lyu, Hassan Gharibi, Susanna L Lundström, Ákos Végvári, Massimiliano Gaetani, Roman A. Zubarev

**Affiliations:** Division of Chemistry I, Department of Medical Biochemistry and Biophysics, Karolinska Institutet, Stockholm, Sweden; Chemical Proteomics Unit, Science for Life Laboratory (SciLifeLab), Stockholm, Sweden; Chemical Proteomics, Swedish National Infrastructure for Biological Mass Spectrometry (BioMS), Stockholm, Sweden; Department of Microbiology, Tumor and Cell Biology, Karolinska Institutet, 17177, Stockholm, Sweden; HDXperts AB, Danderyd, Sweden

## Abstract

Here we present a high-throughput virtual top-down proteomics approach that restores the molecular weight (MW) information in shotgun proteomics, and demonstrate its utility in studying proteolytic events in programmed cell death. With Gel-Assisted Proteome Position Integral Shift (GAPPIS), we quantified over 7000 proteins in staurosporine-induced apoptotic HeLa cells and identified 84 proteins exhibiting in a statistically significant manner at least two of the following features: 1) a negative MW shift; 2) an elevated ratio in a pair of a semi-tryptic and tryptic peptide, 3) a negative shift in the standard deviation of MW estimated for different peptides, and 4) a negative shift in skewness of the same data. Of these proteins, 58 molecules were novel caspase 3 substrates. Further analysis identified the preferred cleavage sites consistent with the known caspase cleavages after the DXXD motif. As a powerful tool for high-throughput MW analysis simultaneously with the conventional expression analysis, GAPPIS assay can prove useful in studying a broad range of biological processes involving proteolytic events.

---

Historically, proteomics was born as a gel-based analysis^1–5^, relying primarily on protein separation by one-dimensional gel electrophoresis (1D-PAGE) and two-dimensional gel electrophoresis (2D-PAGE). 1D-PAGE, also known as Sodium Dodecyl Sulfate-Polyacrylamide Gel Electrophoresis (SDS-PAGE), involved the separation of proteins in a gel matrix based on their molecular weight (MW)^6,7^. Before being loaded onto a polyacrylamide gel, proteins are denatured by SDS and reduced by β-mercaptoethanol. When an electric current is applied, proteins migrate through the gel, with smaller proteins moving faster and farther, and larger ones traveling shorter distances on the gel. After electrophoresis, the gel is typically stained with Coomassie Blue or silver stain to visualize the separated protein bands. The MW scale is calibrated using a separate gel lane with a “ladder” of reference proteins with well-defined MW that exhibit narrow bands. When two conditions are compared, the bands of sample proteins that changed their position or density are excised, digested by trypsin and identified by mass spectrometry (MS).

2D-PAGE is a more advanced and powerful gel-based technique that offers better resolution of proteins and enables differentiation between their proteoforms. 2D-PAGE combines two orthogonal separation dimensions: isoelectric focusing (IEF) in the first dimension and SDS-PAGE in the second dimension^8,9^. The resulting 2D gel image consists of a pattern of spots, each representing a different proteoform. In proteomics analysis, these spots are visualized, and the spots of interest (typically, the ones with altered position or density) are excised for identification by MS, while quantification is based on spot density.

Gel-based proteomics was time-consuming, labor-intensive, and had several other limitations. One of the most serious drawbacks was the inability to identify more than a handful of shifted proteins. Density-based quantification was also a challenge. Despite these limitations, gel-based proteomics played a crucial role in the early days of proteomic research, being instrumental in cancer biomarker discovery^10–12^, neurodegenerative disease research^13–15^, drug development and pharmacology^16,17^, and other important areas.

Subsequently, gel-based methods were replaced by “shotgun” or “bottom-up” proteomics^18,19^, in which the S-S bonds in proteins are first reduced and alkylated, the proteome is digested with trypsin and the peptide mixture undergoes LC-MS/MS analysis. Compared to gel-based proteomics, the shotgun approach has several clear advantages, such as the depth and breadth of analysis, as well as the use of several independently obtained peptide abundances for protein quantification. However, the loss of MW information is an indisputable drawback in the bottom-up approach. This information is particularly important in studying the appearance of abnormal protein fragments, including truncated or cleaved proteins, especially in the context of cell death^20,21^, cancer^22,23^ and other diseases^24,25^.

In the shotgun approach, detection of the protein cleavage is not straightforward, and it is not part of the normal analysis workflow. One indication of such a cleavage is the missing tryptic peptide encompassing the cleavage site. However, the sequence coverage of most proteins in a typical shotgun proteomic analysis is less than 50%, and missing peptides are a common occurrence. Another possible indication is the presence of semi-tryptic peptides. However, semi-tryptic peptides are usually ignored in shotgun proteomics, as their true positive identification involves a significantly higher burden of proof than that of fully tryptic peptides.

As an SDS-PAGE gel would reveal the MW shift in the case of a protein cleavage, a method emerged termed Protein Topography and Migration Analysis Platform (PROTOMAP)^26^ in which the 1D-PAGE gel with the separated proteome is cut into a large number N (20 ≤ N ≤ 100) of narrow bands, with the proteins in each band digested and analyzed separately by LC-MS/MS. The PROTOMAP approach demonstrated its analytical utility by detecting MW shifts in proteins undergoing proteolytic cleavage by caspases and identifying caspase substrates by such shifts^26^. PROTOMAP has also revealed numerous proteolytic events in blood plasma, providing significant coverage of the coagulation degradome^27^. However, to achieve sufficient statistical power, the method required the already large number of gel bands to be analyzed in replicates, resulting in time-consuming sample preparation and a vast number of LC-MS/MS analyses. This ultimately prevented PROTOMAP from becoming a standard method.

Here we present the Gel-Assisted Proteome Position Integral Shift (GAPPIS) assay as a high-throughput alternative to PROTOMAP. GAPPIS derives the MW information on thousands of proteins by analyzing only two gel pieces. To achieve this, each gel lane is cut diagonally along the whole length, providing pieces A and B (Fig. 1a-c). Each protein band becomes therefore split into two parts. The proteins are then extracted from both gel pieces, reduced and alkylated, and then digested. The digests of the extracted proteomes are then labelled by tandem mass tag (TMT) and the TMT-multiplexed samples are analyzed by LC-MS/MS (Fig. 1d-e). It is clear from Fig. 1c that a protein with a higher MW exhibits a higher ratio between the protein abundances in pieces B and A, and vice versa. Thus, the GAPPIS ratio (B/A) will provide an estimate of the protein MW, while the sum (A+B) will reflect the overall protein abundance of each specific protein in the proteome. For more precise MW scale calibration, well-known proteins with minimum post-translational modifications (PTMs) should be used (Fig. 1f).

**Fig. 1.**
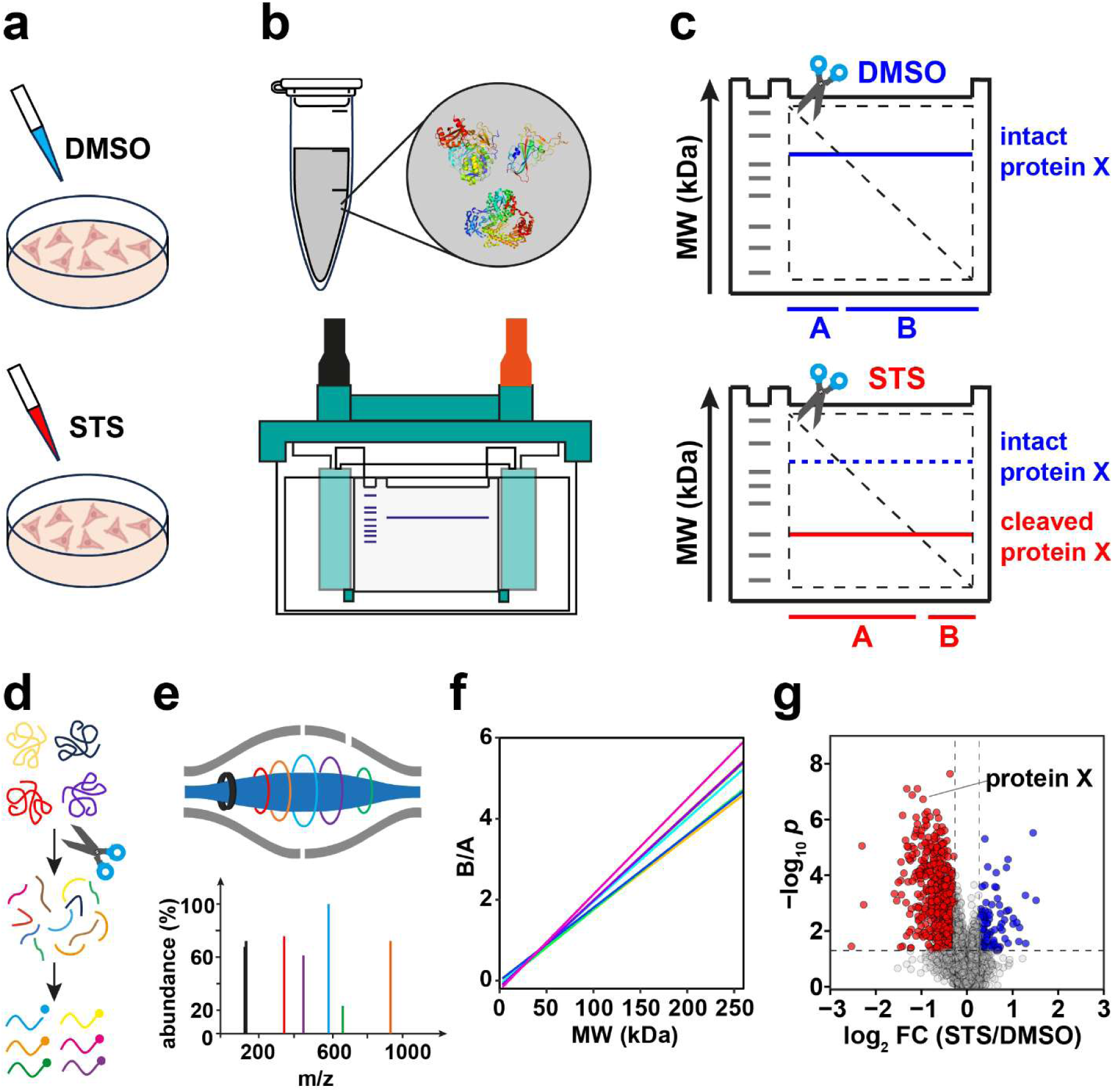
GAPPIS workflow. **(a)** HeLa cells are treated with staurosporine (STS) or DMSO (Control); (b) Cell lysis and SDS-PAGE. (c) Each gel is cut diagonally; the abundance ratio B/A provides the position (MW) of proteins. A cleavage in protein X results in a decrease in its MW and B/A ratio. (d) Gel pieces are digested in-gel, then peptides are extracted and TMT-labelled, followed by fractionation. (e) LC-MS/MS analysis. (f) B/A ratio is converted to MW, providing MW scale calibration using selected proteins and MW estimation for all other proteins. (g) A volcano plot identifies proteins with significant MW shifts.

To test the GAPPIS assay performance, we employed a biological system similar to that in the first PROTOMAP study^26^. Namely, the pan-kinase inhibitor staurosporine (STS) was used to treat HeLa cells, which induced apoptosis and activated caspase 3 proteolytic cleavages. Our goal was to identify the substrates of caspase 3 and compare the GAPPIS results with the three previous works on the subject^26,28,29^. In one study^26^, 261 caspase 3 substrates were identified by PROTOMAP. Prior to that, Lüthi and Martin have compiled the CASBAH database containing all known by then caspase substrates (313 human proteins in total)^28^. Subsequently, Mahrus et al. selectively biotinylated free protein N-termini, performed enrichment of the corresponding N-terminal peptides and identified 282 caspase 3 substrates^29^. The overlap between the caspase substrate lists in these studies ranges between 29% and 42%, which testifies to the need of additional research on the subject. With GAPPIS, we confirmed previously found caspase substrate candidates and found as well as validated new caspase 3 substrates.

## RESULTS

### Establishing MW scale

In the MS2-based analysis, we identified and quantified 7433 proteins based on 103,927 peptides; the corresponding values for the MS3 dataset were 6640 proteins and 69,843 peptides. Fig. 2a demonstrates a strong linear correlation (r = 0.89) observed in the MS3 dataset between protein B/A value and theoretical MW in the range between 3 kDa and 260 kDa. In the MS2 dataset the correlation was also significant, but weaker than in MS3. As the precision of MW determination is defined by the precision of abundance measurements in pieces A and B, it was not surprising that the MS3 approach provided better results than MS2 due to the reduction of the peptide cofragmentation phenomenon that is common in the MS2-based analysis^30^.

**Fig. 2.**
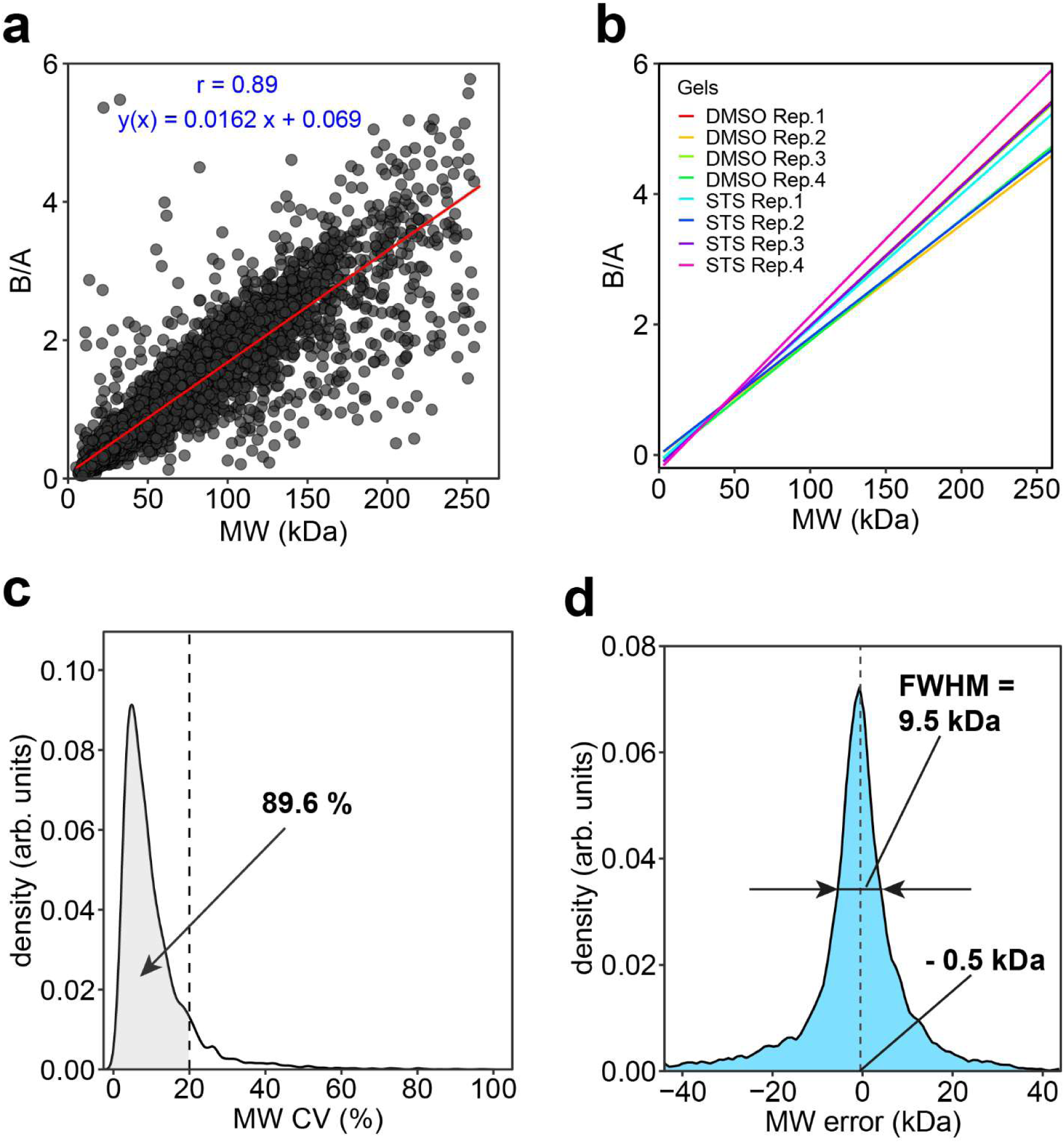
Correlation of the protein GAPPIS ratio (B/A) with MW for MS3 dataset. (a) Correlation of B/A ratios with MW for all 6640 identified proteins. (b) Calibration curves for all 8 gels. (c) CV distribution of the protein B/A-estimated MWs. (d) Error distribution of the B/A-estimated MWs.

To establish the robust MW calibration lines, we selected in the MS3 dataset 641 reference proteins according to the following criteria: a) not to be listed as caspase substrates in either of 3 previous studies^26,28,29^, b) have a sequence coverage of ≥50%, c) be devoid of UniProt-reported post-translational modifications (PTMs), not to be part of mitochondrion, contain transit peptide or being a repressor^31^, d) the B/A values of all peptides should have CV < 30% across the four replicates. The resultant lines are shown in Fig. 2b. All plots demonstrated strong correlations (r > 0.95) between B/A and MW (Fig. S1).

The linear regression between GAPPIS-ratio (B/A) and MW of the reference proteins was used to estimate MW for all other detected proteins. Overall, 89.6% of all proteins exhibited CV between replicate MW estimates of less than 20%, with a peaked frequency of CV at only 4.9% (Fig. 2c). The deviations of the estimated MW from the theoretical value (without any PTMs) form a sharp bell-like distribution (Fig. 2d) centred at -0.5 kDa and with a full width at half maximum (FWHM) of only 9.5 kDa. A similar analysis was conducted on the MS2 dataset, yielding somewhat less accurate results (Supplementary Fig. S2 and S3). Note that in the original PROTOMAP study, the gel was cut into 22 pieces, which for the range of 0-250 kDa corresponds to an MW resolution of ≈11 kDa ^26^. Therefore, the GAPPIS approach is at least as precise as PROTOMAP in MW estimation, despite being based on the analysis of just 2 samples instead of 22.

Since protein MW information is derived in GAPPIS from each peptide independently, it is worth to carefully investigate the distribution of the peptide-level data. For that we selected three proteins with theoretical MW of 24.9, 50.3 and 102.5 kDa, which were not included in the calibration set. For a low-mass protein, the peptide MW-data are not surprisingly localized tightly around the protein’s theoretical MW, while the spread between the peptides increases with MW (Fig. 3a). Besides a longer traveling path on a gel, high-MW proteins are more likely to have multiple isoforms, which explains this result. However, higher-mass proteins also produce on average larger number of peptides, and therefore despite the data spread for individual peptides the center of gravity of the peptides’ positions is rather stable in respect to MW. Analysis showed that the median of the peptide’s MW data is more robust than the average value, likely because medians tend to ignore statistical outliers that more often are false positives. For all three chosen proteins the deviation of the median-derived MWs from the theoretical MW values did not exceed 5 kDa, underscoring the precision of the GAPPIS approach.

**Fig. 3.**
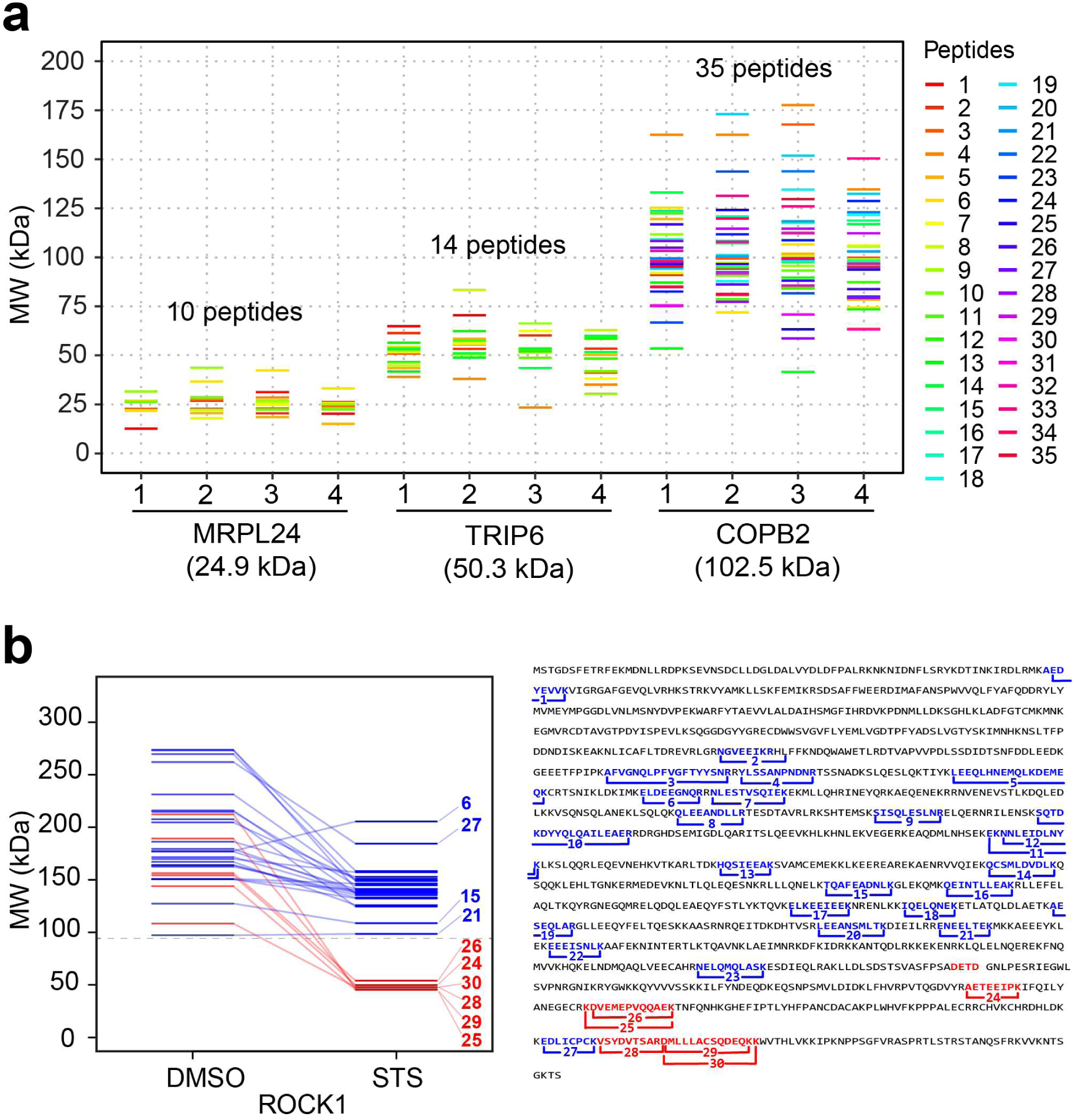
Protein MW estimation from peptide B/A values. (a) A pseudo-gel with three proteins’ MW distributions across 4 replicates in DMSO-treated HeLa cells. (b) Left: A pseudo-gel with peptide-derived MW estimations of the caspase 3 substrate ROCK1 in DMSO-and STS-treated HeLa cells. Right: mapping of the identified peptides on ROCK1 sequence.

It should be noted that the gels are not perfect MW analyzers, as the protein position on the gel is determined by protein mobility that in turn is defined by the protein net charge and its molecular radius. The net charge is determined by the sum of the basic and acidic amino acids in the protein, and the molecular radius - by protein’s tertiary structure. Therefore, systematic deviations of some proteins from the calibration curve should be expected. This deviation is not specific to GAPPIS and affects all gel-based approaches. Yet for practical purposes the molecular mass estimate provided by gels is quite sufficient.

Upon establishing the MW scale, we applied GAPPIS analysis to identification of caspase 3 substrates. To that end we applied four complementary techniques: MW shift, semi-tryptic peptide analysis as well as the novel standard deviation and skewness shift analyses. As the results from these techniques were statistically independent (central moments of a distribution are mathematically orthogonal), an intersection of the proteins supported by at least two techniques was considered to be their validation. We also compared the identified caspase 3 substrates candidates with those reported earlier in literature, and analyzed the preferred motifs of the cleavage sites to verify their origin from caspase activity.

### MW shift analysis

For the protein ROCK1 (theoretical MW 158.2 kDa), a well-established substrate for caspase 3 in STS-treated HeLa cells^32,33^, 30 peptides were identified in the GAPPIS analysis (sequence coverage 32.9 %). The median MW of all these peptides shifted from 173.9±17.9 kDa for untreated sample to 135.2±7.4 kDa for the STS-treated one (p<0.05), immediately identifying this protein as a protease substrate. The known caspase 3 cleavage in ROCK1 produces C-terminal fragment of 130 kDa and N-terminal fragment of 28 kDa in size, with the cleavage site located after the sequence DETD ^34^. In GAPPIS analysis of STS-treated HeLa cells, most of the 24 peptides mapping to the N-terminal cleavage fragment (highlighted in blue in Fig. 3b) are tightly clustering around MW 141±20 kDa, with all 6 C-terminal peptides (highlighted in red) clustering around 48±3 kDa. From that data, the cleavage position could be located between the residues K_1083_ (at the C-terminus of the detected N-terminal peptide NELQMQLASK) and A_1187_ (at the N-terminus of the C-terminal peptide AETEEIPK). We could establish precise location of the cleavage site by semi-tryptic peptide analysis (see below).

On a volcano plot there are many more proteins with a significant decrease in MW upon STS treatment compared to increased MW (102 vs 3, Fig. 4a). This was expected, as the proteolytic activity is the major process in apoptosis. If all the proteins showing increased MW are false positives, the false discovery rate (FDR) of GAPPIS analysis can be estimated as ≍ 3%, which is remarkably low compared to the alternative approaches. A similar analysis of the MS2 dataset identified 155 proteins with a significant decrease in MW (Fig. S4), of which 69 (68%) were the same as with the MS3 approach. There were 25 proteins with increased MW, which estimated the FDR in the MS2 dataset to be 13.8 %. Merging these two datasets led to the identification of 188 potential caspase 3 substrates by negative MW shift. Of these, 66 proteins (35%) overlapped with at least one of the three previous studies, while the expected random overlap is 16 proteins. At the same time, MW of 26 proteins shifted positively; if all these were false positives, FDR in the merged dataset would be estimated around 12%.

**Fig. 4.**
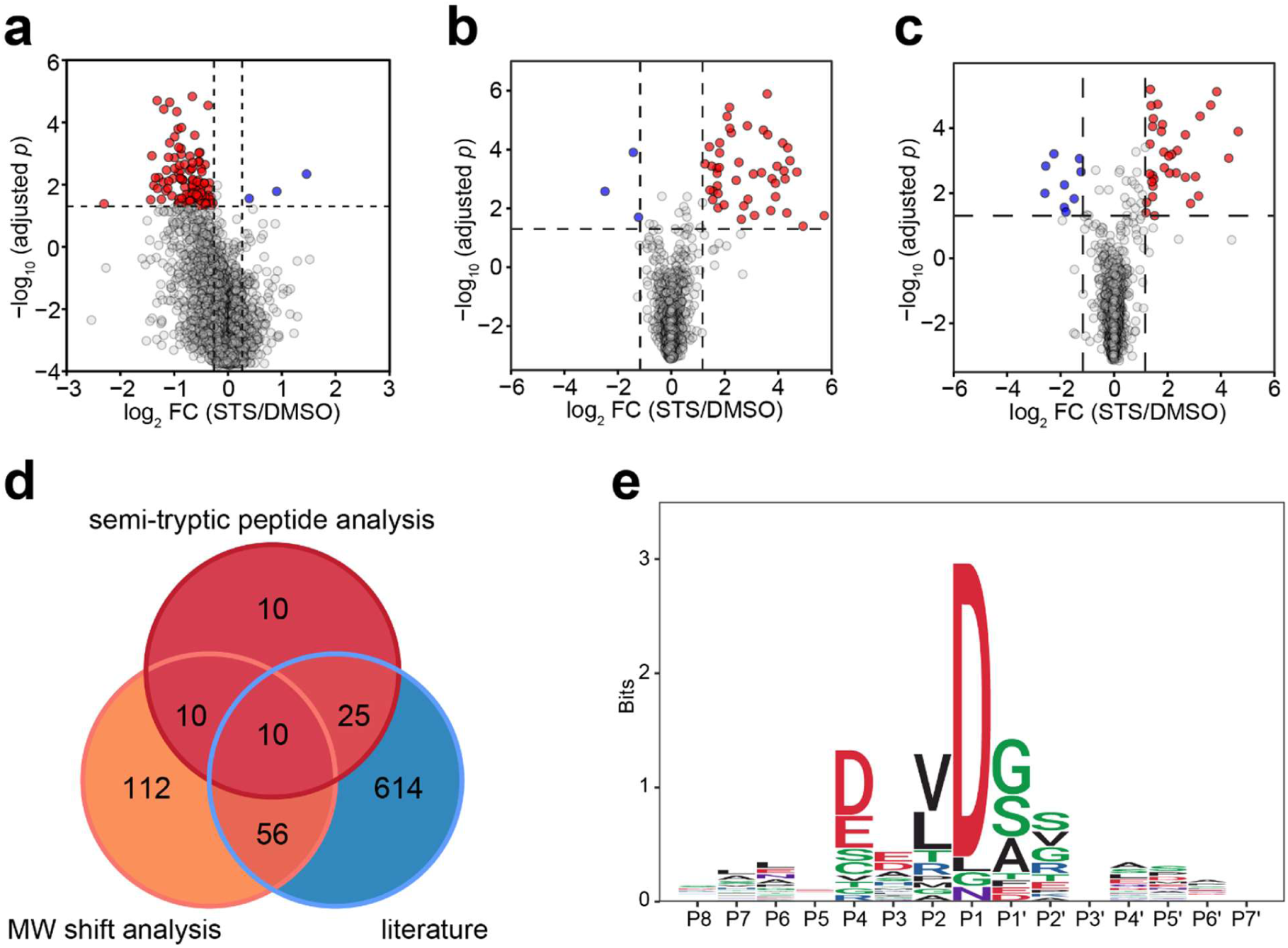
GAPPIS MS3 data identified proteins with significant MW shift. (a) Volcano plot with 102 proteins significantly shifted to lower MW (red) and 3 proteins significantly shifted to higher MW (blue). (b) Volcano plot of semi-tryptic peptides with K/R as the last amino acid residue cells with a significant increase (red) and decrease (blue) in the abundance ratio to fully tryptic peptides STS-treated HeLa cells. (c) Same for semi-tryptic peptides with K/R before the first amino acid residue. (d) Overlap of the GAPPIS-identified caspase 3 substrates with 3 previous studies. (e) Preferred sequence motif for the cleavage sites in 24 semi-tryptic peptides.

### Semi-tryptic peptide analysis

Reliable identification of the cleavage site by semi-tryptic peptides is not an easy task, as the space of possible sequences increases by an order of magnitude compared to fully tryptic peptides, with the FDR increasing proportionally. To enhance confidence in the identified semi-tryptic peptides, we introduced additional requirements as a filter for false discoveries. One such requirement was that the semi-tryptic peptide abundance should increase significantly after STS treatment. Another requirement was that the fully tryptic peptide partially overlapping with a given semi-tryptic peptide should also be present in the dataset. We identified 44 semi-tryptic peptides with a classical tryptic C-terminus that were both paired with their corresponding fully tryptic partner as well as showing a significant increase in their abundance ratio to the tryptic counterpart in the STS-treated HeLa cells compared to DMSO-treated cells (Fig. 4b). A similar analysis was conducted for semi-tryptic peptides with a classical tryptic N-terminus, resulting in the identification of 34 such molecules (Fig. 4c). After merging these results, we identified as caspase 3 substrate candidates a total of 55 proteins in which semi-tryptic peptides increased their abundance after cleavage compared to their fully tryptic counterparts, and 9 proteins with decreased semi-tryptic peptide abundance (if all false positives, FDR ≈14%). Among our caspase 3 substrate candidates 35 proteins were found in literature, giving the record 64% overlap (Fig. 4d), while random match would produce 5 overlapping proteins on average.

Analysis of the preferred cleavage motif was performed using 23 semi-tryptic peptides satisfying the following conditions: a) paired with their fully tryptic counterparts; b) showing significant increase in abundance ratio of semi-tryptic to fully-tryptic peptide in STS-treated HeLa cells; c) stemming from the GAPPIS-identified proteins exhibiting significant negative MW shift in Fig. 4a and Fig. S4. The resultant pattern shown in Fig. 4e revealed that cleavages consistently occurred after the DXXD motif, in line with the known caspase cleavage preference^29,35,36^. This finding further validated the selected proteins as caspase substrates.

### Standard deviation (SD) analysis

As each peptide carries in GAPPIS information on the MW of the protein it belongs to, we used standard deviation (SD), the second central moment of a statistical distribution, as a metric for assessing the dispersion of each protein’s peptide-to-peptide MW estimate. The SD of protein MW exhibited a non-linear upward trend with MW increasing, following the empirical formula *SD* = 0.086*MW*^1.29^ (residual standard error 13 kDa) for DMSO-treatment and *SD* = 0.086*MW*^1.23^ (residual standard error 10 kDa) for STS-treatment (Fig. S5a and S5b). As protein MW decreases after cleavage, we expect a reduction in SD for MW of the substrate proteins. In agreement with this expectation, among the 3384 proteins identified with ≥ 7 peptides (such a peptide number threshold was required for reliable SD estimation), we observed 224 proteins exhibiting a significantly decreased SD after STS treatment compared to DMSO-treated controls (Fig. S5c). At the same time, only 26 proteins displayed a significant increase in SD, which corresponds to a ≈10% FDR if all of the latter proteins were false positives. Out of these 224 proteins, 40 molecules (18%) overlapped with at least one of the three previous studies (Fig. 5), while random match would give 19 proteins on average.

**Fig. 5.**
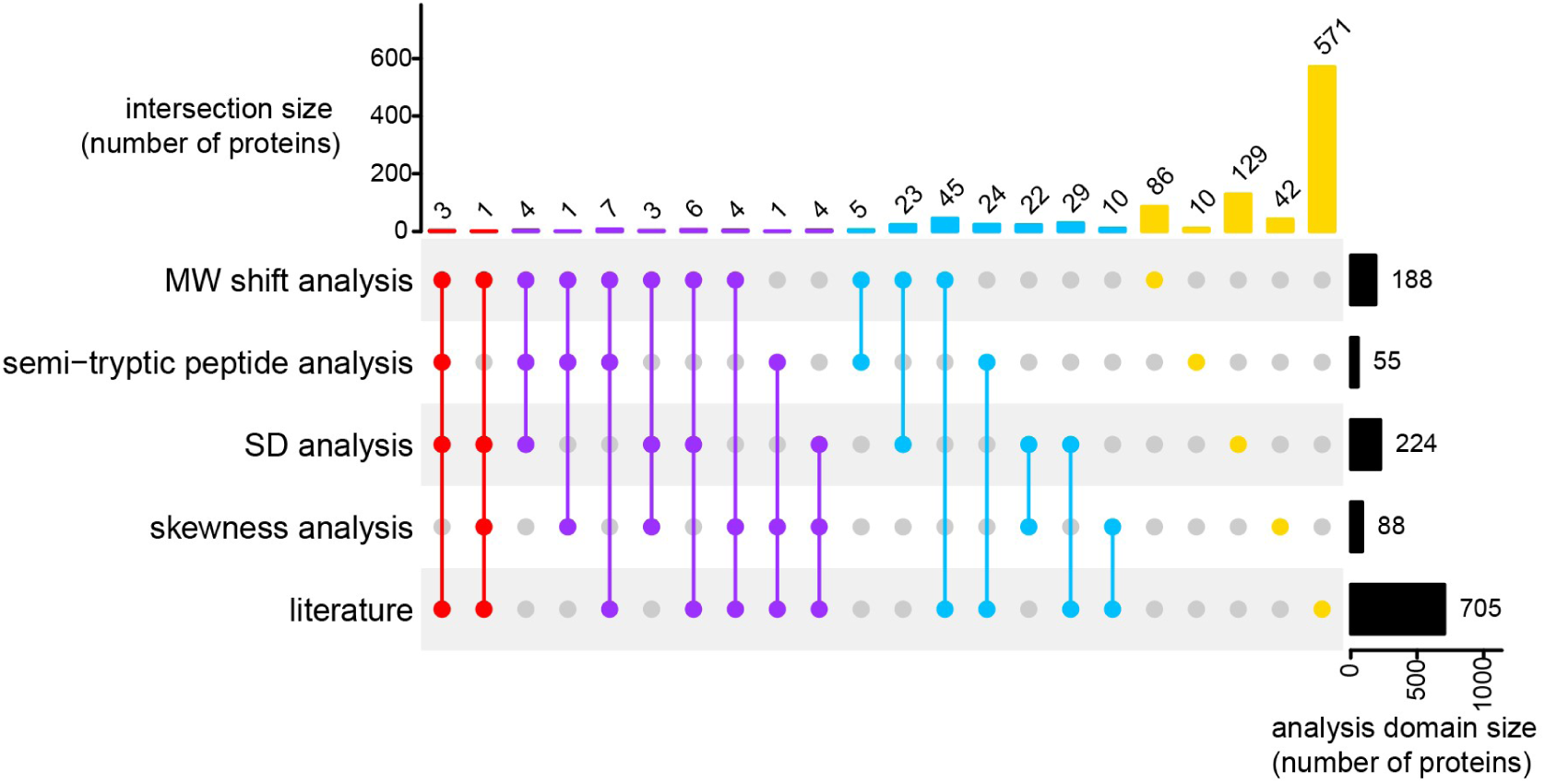
Validation analysis. Overlap between the four independent analysis domains as well as published literature for the GAPPIS-revealed caspase 3 candidate substrates.

### Skewness analysis

Skewness is a third central moment of a statistical distribution, and it is responsible for its asymmetry. Seeking to fully utilize the wealth of information obtained in the GAPPIS approach, we tested whether skewness analysis could be a complementary way of protease substrate identification. The basic assumption was that the peptide MW data distribution of an intact protein should be almost symmetric on the MW scale, being centered around the protein MW, while the presence of two unequal fragments after cleavage would result in an asymmetric distribution skewed on the lower-mass side (negative skewness shift). Consistent with this assumption, among the 1569 proteins identified with ≥ 13 peptides (high threshold needed to obtain precise skewness estimate), we found 88 proteins with significantly decreased skewness and 25 proteins (likely false positives, FDR≈22%) with significantly increased skewness after STS treatment compared to DMSO-treated controls (Fig.S6).

### Validation analysis

As indicated earlier, we considered an overlap between any two of the above complementary analysis domains to be a validation for the caspase 3 candidate substrates. In total, 84 proteins were validated, while the expected random overlap would give less than 8 proteins on average. Of these 84 proteins, 12 proteins were validated by three or more domains, and 26 molecules (31%) were found in literature (Fig. 5, Table S1). Therefore, GAPPIS provided 58 new caspase 3 substrates.

## Conclusions

Here we introduced GAPPIS, a novel shotgun proteomics approach that, being a virtual top-down analysis, brings back the MW information to proteomics. This information, which is orthogonal to both protein abundance and solubility, is obtained from at least an order of magnitude fewer LC-MS/MS analyses than required by the original gel-based PROTOMAP approach, being at least as precise in terms of MW estimation. Clearly, though, should the same protein appear in several bands with distinctly different MWs (e.g., due to multiple proteoforms), the PROTOMAP approach would have had a better chance than GAPPIS of differentiating this from a single-MW situation. However, by applying standard deviation analysis, GAPPIS could still potentially detect an unexpectedly broad distribution of MW data from the peptides belonging to this protein.

The “killer application” of GAPPIS seems to be the same as PROTOMAP, i.e., identification of protease substrates in living cells. There are hundreds of proteases in mammalian cells, implicated in all kinds of biological processes, and the knowledge of their substrates and specificity is important, not least because they represent potential drug targets^37,38^. With GAPPIS, this information may become much more easily available. More importantly, the MW information is encoded in every peptide, both tryptic and semi-tryptic. The wealth of this information is hard to fully appreciate from the standpoint of conventional shotgun proteomics, and in this work, we are just scratching the surface of potential new applications. It is however evident that besides the first central moment (centroid) of the peptide MW distribution, useful information can also be found in the second (standard deviation) and the third (skewness) moments.

As a final comment, the MW information comes in GAPPIS from the precisely measured peptide abundances in gel pieces A and B. The more precise is the abundance measurements, the better is the MW estimation. We found that the MS3-based TMT quantification provides superior performance compared to the easier, more sensitive and much more widely used MS2-based quantification. This finding should encourage further progress in MS instrumentation.

## METHODS

### Cell work and GAPPIS sample preparation

#### Cell culture

Human HeLa cells (ATCC) were grown at 37°C in 5% CO2 using Dulbecco’s Modified Eagle Medium (Lonza, USA) supplemented with 10% FBS superior (Biochrom) and 100 units/mL penicillin/streptomycin (Gibco). Low-number passages (<10) were used for the experiments.

#### Staurosporine treatment and cell lysis

HeLa cells were cultured in 75 cm^2^ flasks for 20 h, subsequently treated with 300 nM staurosporine (STS) or vehicle (DMSO) for 4 h in 4 biological replicates. Cells were washed twice with phosphate buffered saline (PBS) and lysed in M-PER Mammalian Protein Extraction Reagent (Thermo Fisher) supplemented with 1% protease inhibitors (Roche). The cellular lysates were centrifuged at 12,000 rpm for 10 min at 4°C and the soluble fraction was collected. The protein concentration in the lysate was measured using Pierce BCA kit (Thermo Fisher).

#### SDS-PAGE and gel excision

Cell lysate was diluted using M-PER Mammalian Protein Extraction Reagent for all samples to the same protein concentration. The electrophoresis was performed on NuPAGE 4-12% Bis-Tris Mini Protein Gel (Thermo Fisher) with 2 wells in MES Running Buffer under reduced conditions at 150 V for 60 min using the XCell SureLock system (Thermo Fisher). 100 µg of protein were loaded onto the gel for each sample. One STS-treated sample and one DMSO-treated sample were processed in the same tank. Novex Sharp Pre-Stained Protein Standard (Thermo Fisher) was used as a ladder. After electrophoresis the gels were washed and excised diagonally into two parts (A and B), after which each part of the gel was cut into 1 x 1 mm cubes and transferred into 5 mL LoBind tubes. The gel cubes were centrifuged, followed by sequential washes with ammonium bicarbonate/acetonitrile (1:1, v/v) and acetonitrile, each for 10 min. During the acetonitrile incubation, the gel pieces exhibited shrinkage and opacity. Subsequently, all liquid was removed.

#### Reduction and alkylation

25 mM of dithiothreitol (DTT) (Sigma-Aldrich) in 50 mM ammonium bicarbonate was added to fully immerse gel pieces. The samples were then incubated at 56 °C for 30 min. After cooling to room temperature (RT), any remaining liquid was removed. Following this, acetonitrile was added and incubated for 10 min at RT, after which all liquid was removed. Subsequently, 50 mM iodoacetamide (IAA) (Sigma-Aldrich) was added, and the samples were incubated at RT in darkness for 1 h. Then acetonitrile was used to shrink the gel pieces for 10 min with subsequent liquid removal.

#### Trypsin digestion

Each sample was rehydrated at 4 °C for 30 min by adding the same volume of 10 ng/µL trypsin in 50 mM ammonium bicarbonate containing 0.01% ProteaseMAX Surfactant (Promega). During rehydration, the absorbance of the solution was checked, and additional trypsin solution was added to ensure that gel pieces immersed completely. Subsequent incubation was performed for 2 h at 37°C with agitation at 200 rpm. Condensate from the tube walls was collected by centrifugation at 12000 × g for 10 s. The digestion solution with extracted peptides was transferred into new 2 mL tubes, and trypsin was inactivated by adding trifluoroacetic acid (TFA) (Sigma-Aldrich) to a final concentration of 0.5%.

#### TMT-labeling

Samples were cleaned up using Sep-Pak cartridges (Waters) and dried in a DNA 120 SpeedVac Concentrator (Thermo Fisher). The dried peptide samples were reconstituted in 50 mM EPPS buffer (pH 8.5). Acetonitrile (ACN) was added to achieve a final concentration of 30%. Subsequently, TMTpro 16plex reagents (Thermo Fisher) were added at a 5:1 w/w ratio to each sample, followed by a 2 h-incubation at RT. The TMT labelling reaction was quenched by the addition of 0.5% hydroxylamine. All 16 TMT-labelled samples were combined, acidified with TFA, subjected to clean-up using Sep-Pak cartridges (Waters), and dried in a DNA 120 SpeedVac Concentrator (Thermo Fisher).

#### High-pH fractionation

Peptide separation for deeper proteome analysis was carried out using an Ultimate 3000 HPLC system (Thermo Fisher) equipped with a Xbridge Peptide BEH C18 column (25 cm x 2.1 mm, particle size 3.5 μm, pore size 300 Å; Waters) operating at a flow rate of 200 μL/min. Fractionation was achieved through a binary solvent system comprising 20 mM NH4OH in H2O (solvent A) and 20 mM NH4OH in acetonitrile (solvent B). The elution profile was programmed as follows (data for % of solvent B): an initial gradient from 2% to 23% over 42 min, followed by a rapid increase to 52% within 4 min, further elevation to 63% in 2 min, and a subsequent isocratic hold at 63% for 5 min. The elution process was monitored by UV absorbance at 214 nm. A total of 96 fractions, each containing a 100 μL aliquot, were collected. Fractions were subsequently combined in a sequential order (e.g. 1, 25, 49, 73), providing a total of 24 composite fractions.

#### LC-MS/MS analysis

The 24 composite fractions were analyzed by LC-MS/MS using a Orbitrap Fusion Lumos mass spectrometer equipped with an EASY Spray Source and connected to an Ultimate 3000 RSLC nano UPLC system (all - Thermo Fisher). Injected sample fractions were preconcentrated and further desalted online using a PepMap C18 nano trap column (2 cm X 75 μm; particle size, 3 μm; pore size, 100 Å; Thermo Fisher) with a flow rate of 3.5 μL/min for 6 min. Peptide separation was performed using an EASY-Spray C18 reversed-phase nano LC column (Acclaim PepMap RSLC; 50 cm X 75 μm; particle size, 2 μm; pore size, 100 Å; Thermo Fisher) at 55°C and a flow rate of 300 nL/min. Peptides were separated using a binary solvent system consisting of 0.1% (v/v) formic acid (FA), 2% (v/v) acetonitrile (ACN) (solvent A) and 98% ACN (v/v), 0.1% (v/v) FA (solvent B) with a gradient of 3−28% B in 115 min, 28−40% B in 5 min, 40−95% B in 5 min. Subsequently, the analytical column was washed with 95% B for 5 min before re-equilibration with 4% B.

To identify and quantify TMT-labelled peptides, we utilized both the MS2 and SPS MS3 methods. In the MS2 approach, a full MS (MS1) spectrum was first acquired in the Orbitrap analyzer with mass-to-charge ratio (m/z) range from 375 to 1500, nominal resolution 120,000, automated gain control (AGC) target 4 × 10^5^, and maximum injection time of 50 ms. The most abundant peptide ions were automatically selected for subsequent MS/MS (MS2) analysis with a minimum intensity threshold of 2.5 × 10^4^ and a 30 s dynamic exclusion time. MS2 spectra were acquired in the Orbitrap analyzer with the following settings: quadrupole isolation window 1.2 Th, AGC target 1.25 × 10^4^, maximum injection time 120 ms, fragmentation type HCD, normalized collision energy 38%, nominal resolution 50,000, fixed first m/z 110. The number of MS2 spectra acquired per each MS1 spectrum was determined by setting the maximum cycle time for MS1 and MS2 spectra to 3 s (using a top speed mode).

In the MS3 approach, we employed a synchronous precursor selection (SPS) MS3 method in a data-dependent mode. The scan sequence commenced with the acquisition of a full MS spectrum using the Orbitrap analyzer, with m/z range from 375 to 1500, nominal resolution 120,000, AGC target 4 × 10^5^, and maximum injection time of 50 ms. The most abundant peptide ions detected in the full MS spectrum were then selected for MS2 and MS3 analysis, with maximum cycle time set to 5 s (using a top speed mode). MS2 scans were performed in the linear ion trap with quadrupole isolation window 0.7 Th, AGC target 1 × 10^4^, maximum injection time 35 ms, fragmentation type CID, and normalized collision energy 30%. Following the acquisition of each MS2 spectrum, synchronous precursor selection isolated up to 10 most abundant fragment ions with an isolation window 3.0 Th for subsequent MS3 analysis. These fragment ions underwent further fragmentation by HCD with normalized collision energy 55%. Finally, the MS3 spectrum was acquired in the Orbitrap analyzer with a nominal resolution 50,000, AGC target 1 × 10^5^, maximum injection time of 120 ms, and m/z range 110-500.

## Data processing

The raw LC-MS/MS data were analysed by MaxQuant, version 2.2.0.0. The Andromeda search engine was employed to perform MS/MS data matching against the UniProt Human proteome database (version UP000005640_9606, 20607 human sequences). Enzyme specificity was trypsin, with maximum two missed cleavages permitted. When needed, semi-tryptic option was specified for the identification of semi-tryptic peptides in the MS2 dataset. Cysteine carbamidomethylation was set as a fixed modification, while methionine oxidation, N-terminal acetylation, asparagine or glutamine deamidation were used as a variable modification. 1% false discovery rate was used as a filter at both protein and peptide levels. Default settings were employed for all other parameters. Peptide quantification was executed using TMTpro 16plex.

The post-MaxQuant data analysis was the following. The TMT reporter ion abundances for each sample and replicate were normalized by the sum of these abundances. Then for each peptide we calculated the abundance ratio B/A for the two parts of each diagonally-cut gel. The protein B/A values were obtained using the median of the B/A ratios of the peptides associated with a given protein. Following this, we calculated the coefficient of variation (CV) of the B/A ratios between the four replicates of DMSO treatments.

To convert the protein B/A values into MW, we established calibration curves for each gel lane using the proteins chosen as described in the main text. Utilizing the lane-specific calibration curves, we computed the protein MW values for all four replicates of the DMSO/STS-treatment and determined the distribution of the MW errors (deviations from the calibration curve), as well as its parameters, such as the centroid and full width at half maximum (FWHM).

After converting B/A values of all identified proteins to MW using calibration curves, volcano plots were generated by calculating the MW fold change (FC) between STS and DMSO treatments and conducting Student’s t-tests for the two treatment groups. Given the follow-up validation, false discovery rate (FDR) of ≤10% was determined to be acceptable for proteins with significant MW shifts (FDR is the number of proteins with significant MW increase divided by the total number of proteins with both significant MW increase and decrease). The p-value adjustment method was peptide number-based correction: the peptides were ranked in descending order according to the number of peptides attributed to each protein, and p-values were subsequently adjusted by multiplying by their respective ranks.

## Data availability

The data supporting the findings of this study are available in the supplementary information files. The mass spectrometry data have been deposited in the ProteomeXchange Consortium via the PRIDE partner repository (https://www.ebi.ac.uk/pride/) with the dataset identifiers PXD049007.

## Notes

### Competing Interest Statement

The authors have declared no competing interest.

### Summary of Updates

No significant differences between this version and previous version of the manuscript.

