## Supplementary Figures for "Gel-Assisted Proteome Position Integral Shift (GAPPIS) Assay Returns Molecular Weight to Shotgun Proteomics and Identifies Novel Caspase 3 Substrates"

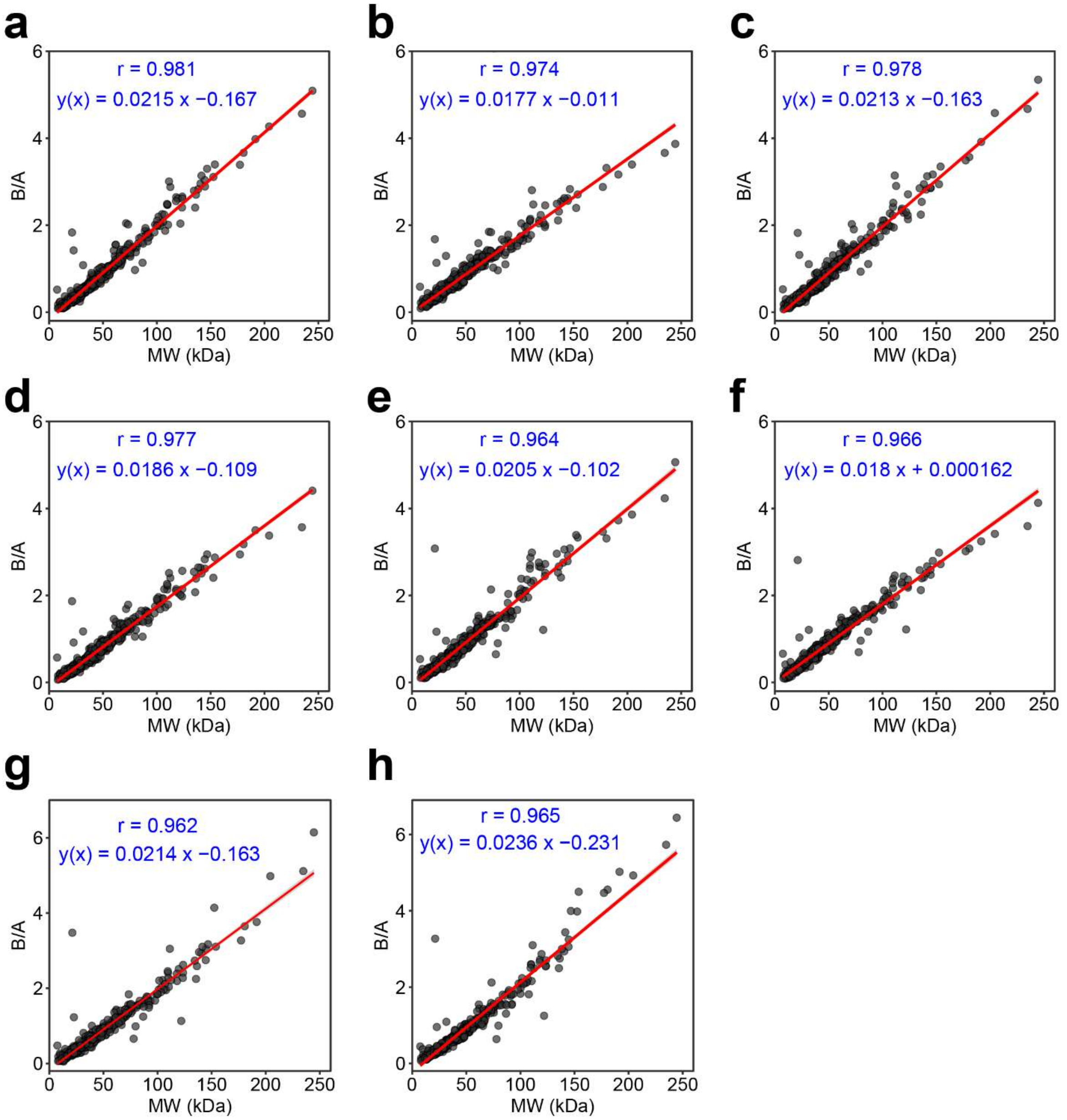


**Fig. S1. Calibration curves converting B/A values to MW for MS3 data.** The reference proteins selected to plot calibration curves have peptides sequence coverage ≥ 50%, do not have reported PTMs and do not overlap with 3 previous studies. (a) to (d) Data calibration curves for the four replicates of DMSO-treated HeLa cells, respectively. (e) to (h) Data calibration curves for four replicates of STS-treated HeLa cells, respectively.

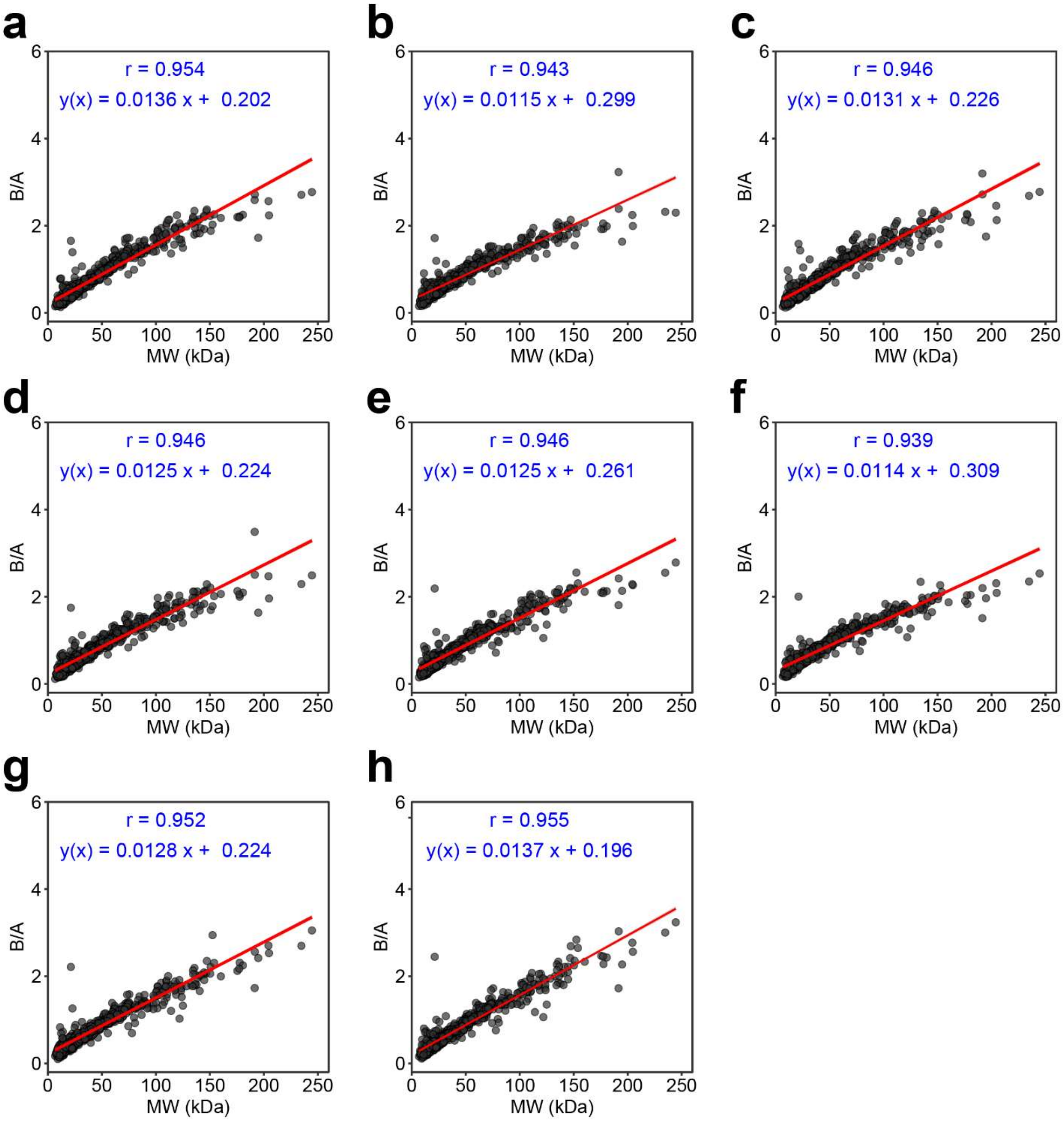


**Fig. S2. Calibration curves converting B/A values to MW for MS2 data.** The reference proteins selected for plotting calibration curves have peptides sequence coverage ≥ 50%, do not have reported PTMs and do not overlap with 3 previous studies. (a) to (d) Data calibration curves for the four replicates of DMSO-treated HeLa cells, respectively. (e) to (h) Data calibration curves for four replicates of STS-treated HeLa cells, respectively.

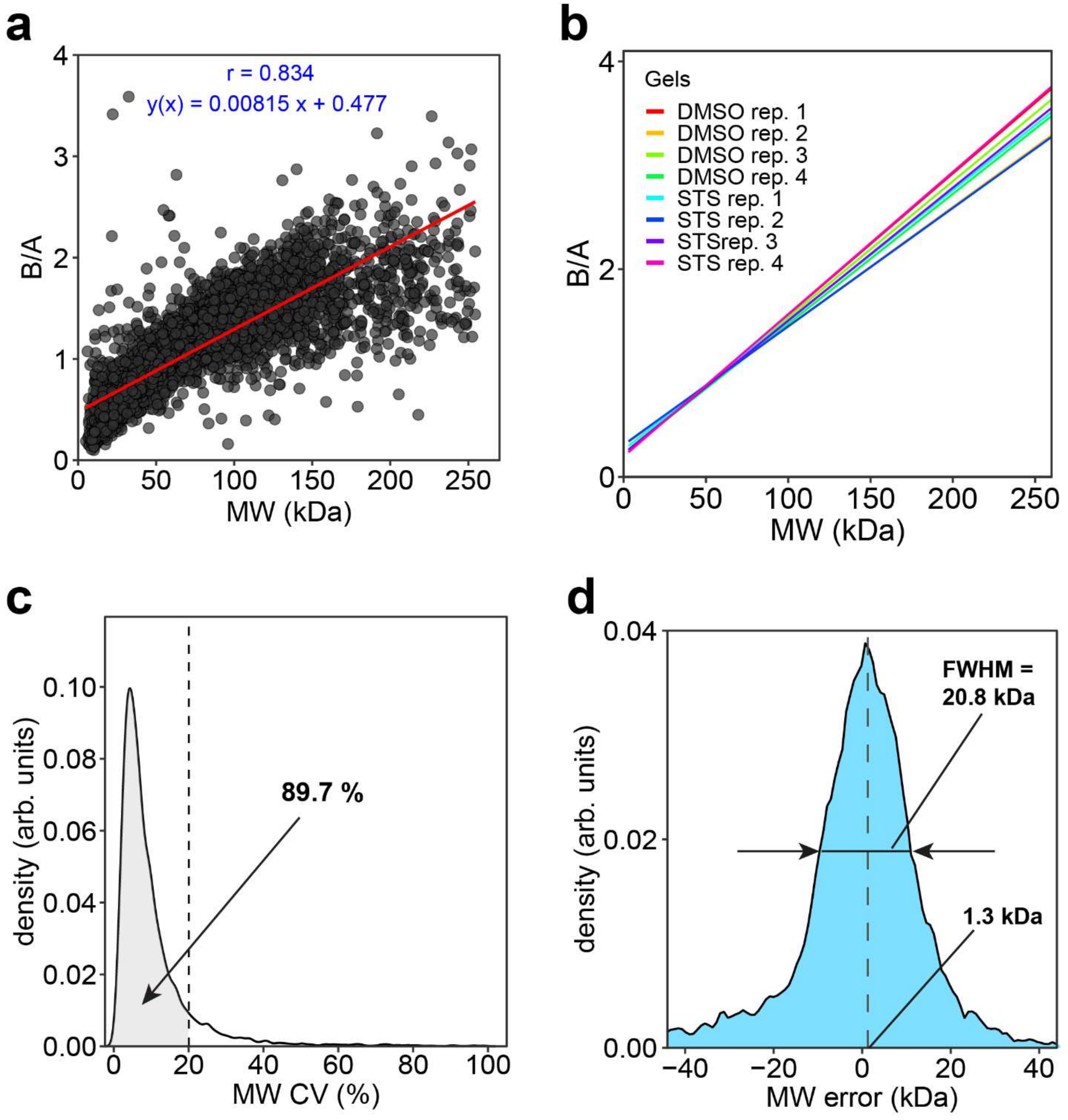


**Fig. S3. Protein MW estimations from peptide B/A ratios for MS2 dataset.** (a) Correlation of B/A ratios with MW for all 7433 proteins. (b) Calibration curves for all 8 gels. (c) CV distribution of B/A-calculated protein MW values. (d) Error distribution of B/A-calculated protein MW values.

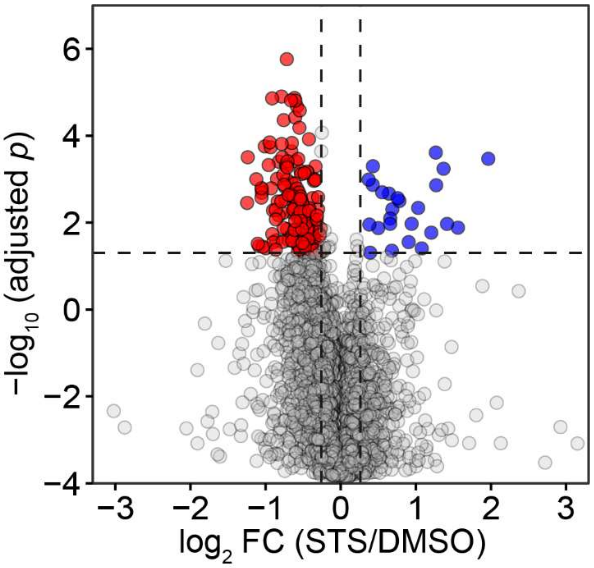


**Fig. S4. Volcano plot of proteins with significant MW shifts in MS2 dataset.** With p values adjusted by peptides number-based multiple hypothesis correction, volcano plot shows 155 proteins significantly shift to lower MW (red) while 25 proteins significantly shift to higher MW (blue).

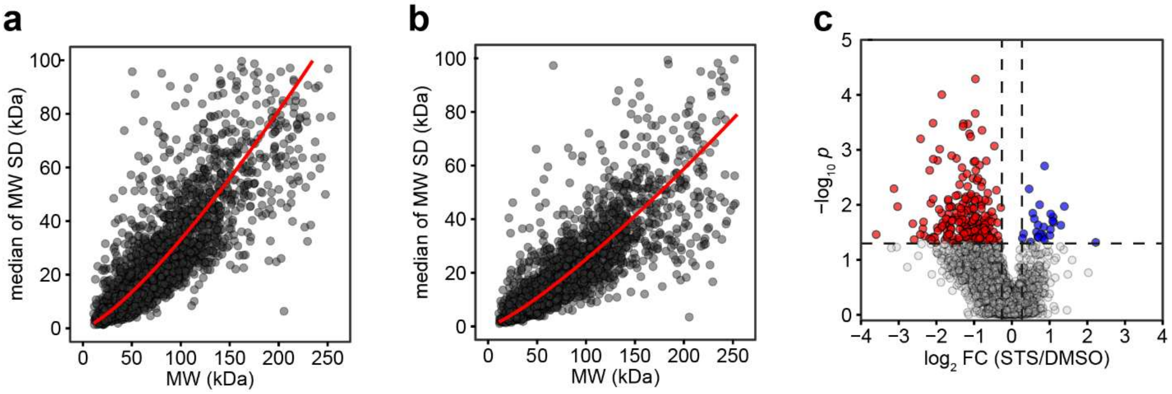


**Fig. S5. Standard deviation (SD) analysis for identifying caspase substrate candidates.** (a) The empirical formula $SD = 0.086MW^{1.29}$ is fitted to the SD-MW plot for DMSO-treatment, with residual standard error of 13 kDa. (b) The empirical formula $SD = 0.086MW^{1.23}$ is fitted to the SD-MW plot with residual standard error 10 kDa for STS-treatment. (c) Volcano plot for 3384 proteins with ≥ 7 peptides from STS-treated HeLa cells with proteins showing significantly decreased (red) and increased (blue) SD.

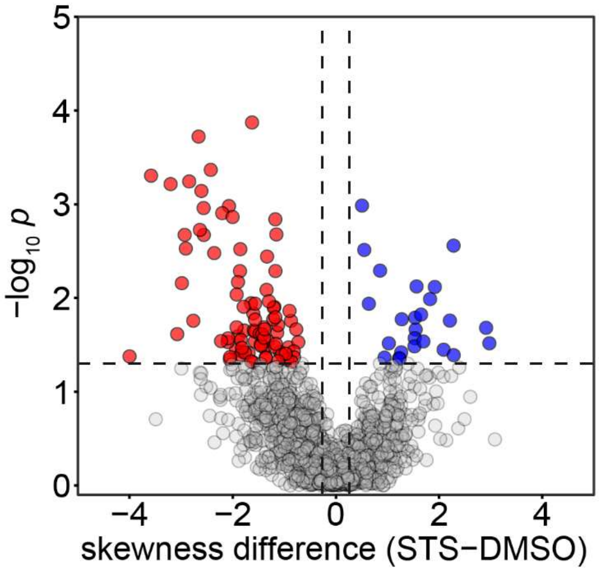


**Fig. S6. Skewness analysis for identifying caspase substrate candidates.** Volcano plot for 1569 proteins with ≥ 13 peptides showing in STS-treated HeLa cells proteins with significantly decreased (red) and increased (blue) skewness.
